# Natural and engineered mediators of desiccation tolerance stabilize Human Blood Clotting Factor VIII in a dry state

**DOI:** 10.1101/2022.11.28.518276

**Authors:** Maxwell H. Packebush, Silvia Sanchez-Martinez, Sourav Biswas, KC Shraddha, Kenny Nguyen, Thomas C. Boothby

## Abstract

Biologics, pharmaceuticals containing or derived from living organisms, such as vaccines, antibodies, stem cells, blood, and blood products are a cornerstone of modern medicine. However, nearly all biologics have a major deficiency: they are inherently unstable, requiring storage under constant cold conditions. The so-called ‘cold-chain’, while effective, represents a serious economic and logistical hurdle for deploying biologics in remote, underdeveloped, or austere settings where access to cold-chain infrastructure ranging from refrigerators and freezers to stable electricity is limited. To address this issue, we explore the possibility of using anhydrobiosis, the ability of organisms such as tardigrades to enter a reversible state of suspended animation brought on by extreme drying, as a jumping off point in the development of dry storage technology that would allow biologics to be kept in a desiccated state under ambient or even elevated temperatures. Here we examine the ability of different protein and sugar-based mediators of anhydrobiosis derived from tardigrades and other anhydrobiotic organisms to stabilize Human Blood Clotting Factor VIII under repeated dehydration/rehydration cycles, thermal stress, and long-term dry storage conditions. We find that while both protein and sugar-based protectants can stabilize the biologic pharmaceutical Human Blood Clotting Factor FVIII under all these conditions, protein-based mediators offer more accessible avenues for engineering and thus tuning of protective function. Using classic protein engineering approaches, we fine tune the biophysical properties of a protein-based mediator of anhydrobiosis derived from a tardigrade, CAHS D. Modulating the ability of CAHS D to form hydrogels made the protein better or worse at providing protection to Human Blood Clotting Factor VIII under different conditions. This study demonstrates the effectiveness of tardigrade CAHS proteins and other mediators of desiccation tolerance at preserving the function of a biologic without the need for the cold-chain. In addition, our study demonstrates that engineering approaches can tune natural products to serve specific protective functions, such as coping with desiccation cycling *versus* thermal stress. Ultimately, these findings provide a proof of principle that our reliance on the cold-chain to stabilize life-saving pharmaceuticals can be broken using natural and engineered mediators of desiccation tolerance.

## Introduction

### Biologics are a cornerstone of modern medicine

The past two decades have seen an explosion in the development and use of ‘biologics,’ drugs derived from or containing components of living organisms. This includes vaccines, protein and nucleic acid-based pharmaceuticals, allergens, anti-venoms, blood, blood components, and cell-based therapeutics. The increase in biologic development has been mirrored by their increased use in our healthcare systems, for example, between 1999 and 2006 there was an increase from 3% of patients in the United States being prescribed a biologic to 26%^1^. Additionally, by the end of 2020, biologics accounted for over 1/4th of the world’s pharmaceuticals^2^.

### The fragility of biologics is their major drawback

While biologics have proven effective in combating numerous diseases they have a major drawback, they are inherently unstable^3^. The breakdown, aggregation, and/or modification of biologics due to improper storage or transportation is an enormous economic burden. In 2019, ~35% of all vaccines in developing countries had to be discarded due to improper maintenance, leading to multiple countries reporting losses greater than $1,500,000 for compromised vaccines alone^3^. Perhaps more troubling, not only does the breakdown of biologics lead to reduced efficacy and economic losses, but in many cases loss of biologic integrity can result in ‘degradation products’ with detrimental, instead of health-inducing, effects^4^. Because of their inherent instability, the economic loss associated with their breakdown, and the risk of harmful effects being induced by breakdown products, it is essential that we employ reliable, effective, and economical means of stabilization for these life-saving medicines.

### The cold-chain is not a universally reliable, effective, and economical means of stabilizing biologics

Currently, the most widespread method of biologic stabilization is cold storage. Cold storage of biologics uses what is known as the ‘cold-chain’, a system of refrigerators and freezers used during the production, transportation, and storage of a biologic to help maintain its viability. Currently, ~3/4th of all biologic drugs, including essentially all vaccines, ~15% of all small-molecules, as well as myriad biological samples and diagnostic tools rely on the cold-chain for stabilization^5^. While the cost of the global cold-chain is staggering and steadily growing, $17.2 billion in 2020 and forecasted at $21.3 billion in 2024, under ideal circumstances cold stabilization can be effective^5^. However, in remote or developing parts of the world, purchasing and maintaining the necessary infrastructure such as freezers, electrical systems, and backup generators needed for the cold-chain to work seamlessly can be close to impossible. A prime example of the economic burden of the cold-chain is the fact that in developing countries between 40-90% of the cost of a vaccination program comes from the need to keep vaccines cold^6^. This means that for underdeveloped countries, where it is estimated that ~50% of all healthcare facilities completely lack any electrical infrastructure, with only ~10% having access to reliable electricity, the cold-chain is neither efficient, reliable, or economically viable^7^.

### Dry preservation is an alternative to the cold-chain

The cold-chain is currently essential for the stabilization of most biologics. This is because above certain temperatures (e.g., ~8 °C for most vaccines) molecular dynamics and rearrangements are accelerated leading to hastened breakdown of these materials^5^. At low temperatures (e.g., ~2 °C for most vaccines) water begins freezing leading to the formation of ice crystals that can irreparably damage sensitive biologics^5^. Therefore, an effective means of stabilization should: i.) prevent or reduce molecular motion and ii.) minimize crystallization. However, reduced molecular motion and inhibition of crystallization are not unique to cold-stabilization, and there are myriad examples of biological stabilization in nature that do not rely on cold temperatures^8–10^. An extreme example of such stabilization is anhydrobiosis (from Greek meaning ‘life without water’) which is a term used to describe the ability of some organisms to lose essentially all the water inside their cells, enter a state of suspended animation, and remain viable in that dry, ametabolic state for years and sometimes decades^11–13^. Examples of organisms that can undergo anhydrobiosis are spread across the biological kingdoms, including life stages of nearly every land plant (e.g., seeds, spores, pollen), many bacteria and archaea, fungi, protists, and even some animals - notably tardigrades, brine shrimp, rotifers and some nematodes^8,9,14,15^.

An overarching theme among how these desiccation-tolerant organisms stabilize their sensitive biological macromolecules is their reliance on special protectants, historically recognized as sugars such as trehalose, that promote intracellular vitrification or glass formation^12,13,16^. Vitrification is thought to be protective to these organisms in two ways. First, by filling their cells with vitrifying protectants, these organisms increase intracellular viscosity to the point where detrimental effects of drying, such as protein unfolding and membrane fusion as well as normal biological turnover are slowed to such that they essentially stop. Secondly, because these protectants form vitrified solids, which have an amorphous molecular structure, they do not form potentially damaging crystals^12,13,16^. This means that drying induced vitrification observed in nature accomplishes the two main tasks of the cold-chain, reducing molecular motion and minimizing crystallization, but without the need for cold-temperatures.

A long-term promise of the desiccation tolerance field has been the application of natural mediators of anhydrobiosis to the stabilization of biologics in a dry state, so called xeroprotection. However, while natural products often work well in their host organism, they do not always work in heterologous, *ex vivo*, or *in vitro* settings. Furthermore, there are often not obvious means for modification or engineering of these natural products to make them more amenable for biologic stabilization as many of them are sugars or small molecules without clear or empirically determined mechanisms of action.

To address this major obstacle, and to begin to build a foundation for the development of technology to allow for the dry stabilization of diverse biologics, we have searched for proteinbased mediators of anhydrobiosis since these would allow us to use well-established analytical and protein engineering approaches to i.) understand the mechanisms underlying protein-mediated anhydrobiosis and ii.) modify these natural products to tune their properties and functions allowing for their application in the dry preservation of biologics.

To this end, we turn to a recently discovered family of proteins, found exclusively in desiccation tolerant tardigrades, that are both necessary and in some cases sufficient to confer desiccation tolerance *in vivo, ex vivo*, and *in vitro^11,12,17^–^19^*. These proteins, termed Cytoplasmic Abundant Heat Soluble (CAHS) proteins, are expressed at high levels and polymerize to form gels that slow diffusion, preventing desiccation-sensitive proteins from becoming nonfunctional during desiccation and upon rehydration^12,17,19^. Ultimately, as these protective gels dehydrate, they remain amorphous, forming a non-crystalline, solidified matrix in which desiccation-sensitive proteins are presumably embedded and protected from the harmful effects of drying^12,20^. As such, CAHS proteins provide a unique opportunity on which to build a foundation for the development of technology for the dry stabilization of biologics, in that they are amenable to well-established protein engineering approaches and perform the main tasks of cold-storage: i.) reducing molecular motion and ii.) forming non-crystalline solids.

Here we examine the ability of a model CAHS protein originating from the tardigrade *Hypsibius exemplaris*, CAHS D, and other known mediators of desiccation tolerance to stabilize an important, but labile biologic, Human Blood Clotting Factor VIII (FVIII) in a dry state. We selected FVIII because it is an essential component in the intrinsic blood clotting pathway^21^. FVIII alone or in conjunction with other additives, has many therapeutic applications ranging from treatment of genetic diseases (e.g., hemophilia A) to instances of extreme physical trauma^22^. FVIII, like many biologics, is sensitive, with the best current practices requiring cold-storage to maintain its stability and functionality^23^. Thus, developing dry preservation methods for FVIII will assist in deployment of this life saving biologic to remote (*e.g*. space mission), underdeveloped (*e.g*. clinics that lack electricity), and austier (*e.g*. battlefield or areas experiencing natural disaster) settings.

Here we find that classic sugar-based mediators of desiccation tolerance provide some protection to FVIII during repeated desiccation cycles in a concentration dependent fashion. Furthermore, CAHS D provides protection to FVIII during repeated desiccation cycles at concentrations below its gelation point. We test whether gelation inhibits protection of FVIII by utilizing two CAHS D variants, one of which gels while the other does not. Protection, or lack thereof, correlates with the gelled state of these variants, where our gelling variant (2X Linker) protected FVIII below but not above its gel point. Conversely, our non-gelling variant (Linker Region) protected FVIII at a much wider range of concentrations. Supporting these observations, we find that the protective effect of AavLEA1, a protective protein derived from a nematode worm, which does not gel, provides robust protection to FVIII across a wide range of concentrations. However, robust protein-based protection does not extend to all proteins tested, demonstrating that there are unique sequence features and biophysical/biochemical properties that drive protection during repeated desiccation cycles.

Similar to repeated desiccation cycles, we find the sugar-based protectants confer protection to FVIII during thermal stress in a dry state. Interestingly, the strong gelation capacity of the 2X Linker variant, while inhibitory to protection during repeated desiccation cycles, is protective under thermal stress, suggesting that different engineered biophysical and material properties can tune the proteins protective capacity for FVIII under different storage/stress conditions.

Finally, we examine the ability of one of our best performing mediators (CAHS D Linker Region) to extend the shelf-life of dry FVIII. We find that while in a hydrated state CAHS D Linker does not help with shelf-life, but under dry conditions, the stability of FVIII coincubated with the Linker variant was unchanged over 10 weeks, while FVIII dried without excipients degraded significantly over that period of time, as did hydrated samples.

Our results indicate that both natural and engineered mediators of desiccation tolerance can stabilize biologics, such as Human Blood Clotting Factor FVIII, during repeated desiccation cycles and under thermal stress. Furthermore, protein engineering can be used to fine tune the protective capacity of natural mediators of desiccation tolerance to optimize them for different storage conditions. Finally, simple coincubation with mediators of desiccation tolerance can provide long-term, cold-chain independent, storage solutions to the FVIII biologic.

## Results and Discussion

### Classic sugar-based mediators of desiccation tolerance stabilize FVIII during repeated cycles of drying and rehydration

The ability of non-reducing disaccharides, such as sucrose and trehalose, to stabilize client biomolecules in a dry state has long been established^17,19,24–26^. Because of this, we selected trehalose and sucrose for our initial investigations into whether FVIII could be stabilized and stored in a dry state without refrigeration. To start, we subjected samples to 6 repeated cycles of desiccation and rehydration. This particular stress simulates partial rehydration and dehydration implicit with long term storage and transit where humidity is not absolutely controlled such as in clinical settings in remote or developing parts of the world. This desiccation regime also explores the possibility that a dry pharmaceutical could be rehydrated, used, desiccated, and reused in the future.

To determine the functionality of FVIII after repeated desiccation and rehydration cycles, we used FVIII deficient human blood plasma (FVIII DHBP) that, without supplementation of FVIII biologic, clots slowly because it lacks proper intrinsic pathway activation (Fig. 1). Function of FVIII was evaluated before and after desiccation by mixing it with FVIII DHBP and measuring 50% clotting time. To obtain a baseline for healthy blood 50% clotting time, we mixed unperturbed FVIII with FVIII DHBP and found that this supplemented plasma had a 50% clotting time of ~150 seconds in our *in vitro* system (Fig. 2A&B, red bar).

**Figure 1.**
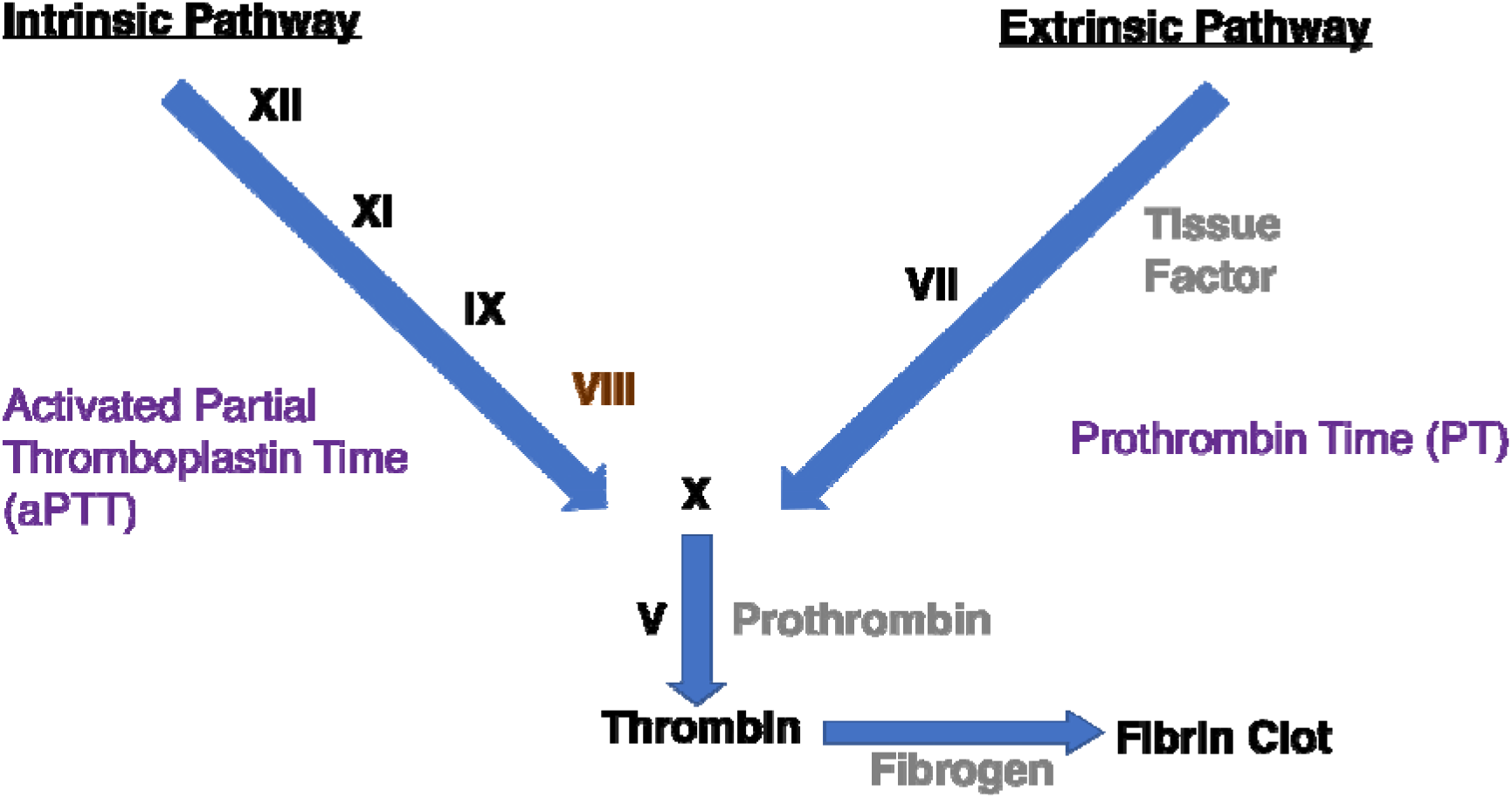
The human blood clotting cascade. The clotting cascade of human blood plasma follows two prominent pathways; intrinsic, measured by Activated Partial Thromboplastin Time (aPTT) and extrinsic, measured by Prothrombin Time (PT). In order to activate the intrinsic pathway, Human Blood Clotting Factor XII (FXII) acts as the first protein in a cascade of clotting factor activation. FXII activates FXI which activates FIX which finally activates FVIII. FVIII subsequently binds to and activates - FX. In order to activate the extrinsic pathway, FVII forms a complex with Tissue Factor (TF), activating FX. After activation of FX, both coagulation pathways converge. FX forms a complex with FV, converting prothrombin into thrombin. Thrombin then converts fibrinogen into fibrin, in turn creating a fibrin clot. Human plasma deficient in Factor VIII (highlighted in red) is unable to clot properly through the intrinsic pathway, unless supplemented with this factor, and thus clots more slowly. Adapted from Zaragoza and Espinoza-Villafuerte, 2017^39^.

**Figure 2.**
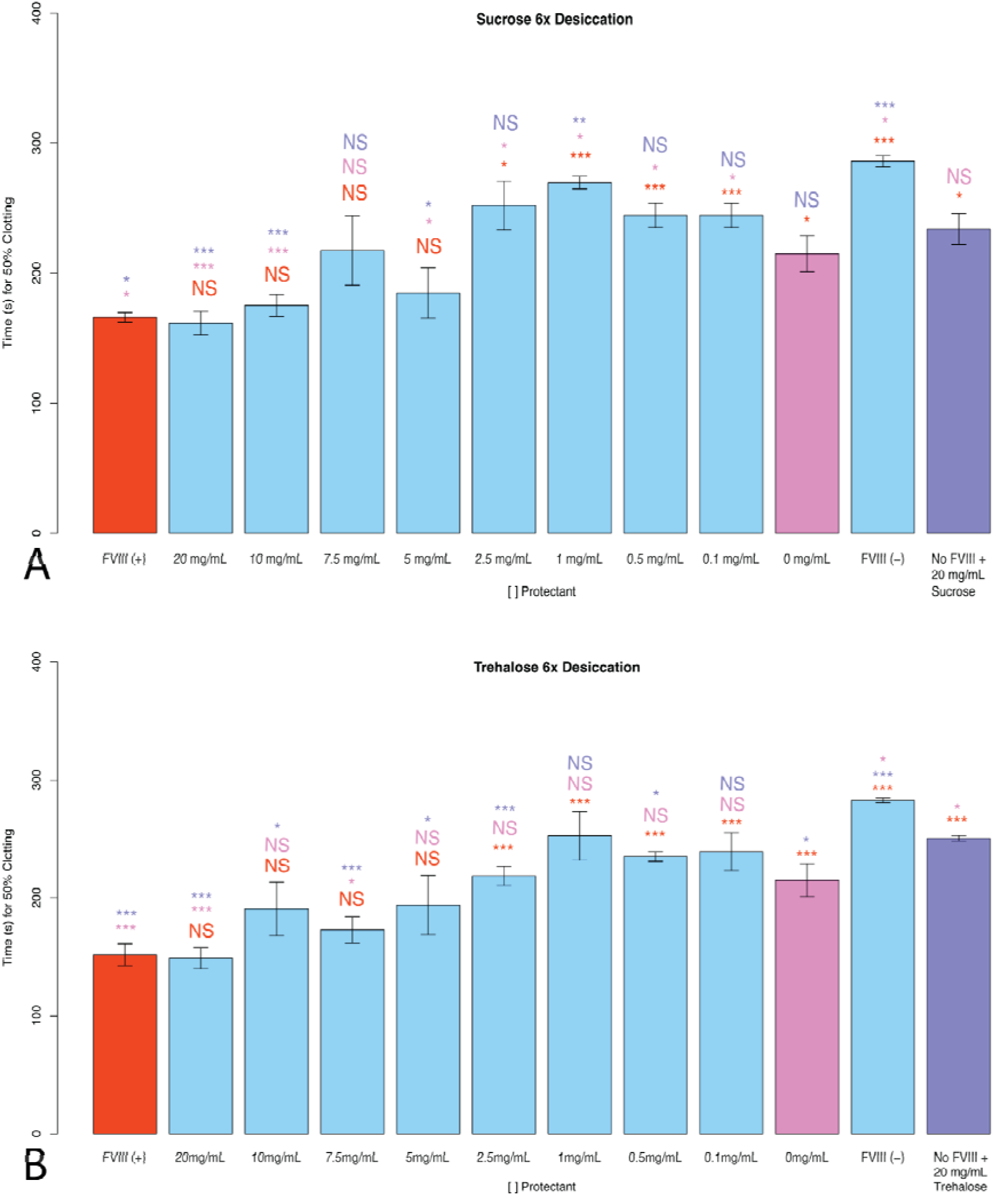
Non-reducing disaccharides stabilize FVIII under repeated desiccation cycles. Histogram showing the 50% clotting times for FVIII coincubated with (A) sucrose or (B) trehalose. FVIII (+) represents FVIII deficient human blood plasma treated with FVIII that has not been desiccated. All red statistical notations above sample bars represent statistical comparison with FVIII (+). “X mg/mL” notations beneath sample bars correspond to the concentrations of non-reducing disaccharide mixed with FVIII before desiccation. All pink statistical notations above sample bars represent statistical comparison with “0 mg/mL of sugars.” FVIII (-) represents FVIII deficient human blood plasma with no supplemented FVIII. “No FVIII + X mg/mL Sugar” represents FVIII deficient human blood plasma treated with the indicated concentration of non-reducing disaccharide, but no FVIII. All purple notations above sample bars represent statistical comparison with “No FVIII + X mg/mL Sugar.” Notations above sample bars represent statistical comparisons by 2-tailed T-test. Error bars represent bidirectional standard deviation. P value > 0.05 = NS, P < 0.05 = *, P < 0.01 = **, P < 0.005 = ***

To determine a negative control baseline, the 50% clotting time of FVIII DHBP without FVIII addition was measured. FVIII DHBP alone had a 50% clotting time of ~200 seconds (Fig. 2A&B), which is significantly slower than FVIII DHBP treated with healthy FVIII (Fig. 2A&B).

To assess the effect of trehalose or sucrose on the 50% clotting time of FVIII DHBP, we treated samples of this deficient plasma with only sucrose or trehalose, and no FVIII. In these cases, the 50% clotting time sped up significantly, but was still not as fast as plasma supplemented with FVIII (Fig. 2A&B, purple bar).

We then subjected the FVIII to 6X desiccation/rehydration stress with either sucrose or trehalose at various concentrations and used these stressed FVIII samples to supplement FVIII DHBP. Desiccation of FVIII without trehalose or sucrose (0 mg/mL) resulted in a clotting factor with compromised clotting ability (Fig. 2A&B). FVIII function was not protected by the addition of 0.1-2.5 mg/mL of sucrose prior to desiccation (Fig. 2A). When dried with higher concentrations of sucrose (5, 10, and 20 mg/mL), the function of FVIII was partially protected or fully preserved (Fig. 2A). A similar concentration dependent trend was observed when trehalose was added to FVIII before desiccation (Fig. 2B).

This data demonstrates that desiccation without excipients compromises the clotting function of FVIII. Furthermore, although sucrose and trehalose themselves speed up clotting in FVIII DHBP, it is apparent that they also protect the function of FVIII during repeated cycles of dehydration and rehydration, since FVIII with 20 mg/mL of sucrose or trehalose caused statistically faster 50% clotting time than FVIII DHBP treated with 20 mg/mL of sucrose or trehalose alone (Fig. 2A&B).

### CAHS D and engineered CAHS D variants stabilize FVIII during repeated cycles of drying and rehydration

Beyond disaccharide xeroprotectants such as trehalose and sucrose, certain intrinsically disordered proteins (IDPs) are also known to confer protection to biological material during desiccation^15^. Proteins represent attractive potential xeroprotectants due to the relative ease with which their behaviors and functions can be engineered. CAHS D is an IDP derived from tardigrades which has the ability to protect essential biomolecules under desiccation stress *in vivo* and *in vitro^11,12,15,17,19^*. These observations make this protein an attractive potential excipient for biologic pharmaceuticals. To assess whether desiccation-related IDPs can be used to stabilize FVIII, we purified and coincubated CAHS D with FVIII prior to 6X desiccation cycles.

Before testing CAHS D, we established a baseline for protection of FVIII with a protein unrelated to desiccation tolerance. In this case, we treated FVIII with a concentration range of lysozyme originating from hen egg-white. Lysozyme is a well characterized and highly structured protein with no link to desiccation tolerance^27^. As expected, after 6 desiccation cycles, FVIII treated with lysozyme at any concentration showed at best modest functionality, and at high concentrations lysozyme was antagonistic rather than protective (Fig. 3A). This is in contrast to trehalose and sucrose which provide not only partial but also complete protection of FVIII at some concentrations (2A&B) - as well as some concentrations of CAHS D (see below).

**Figure 3.**
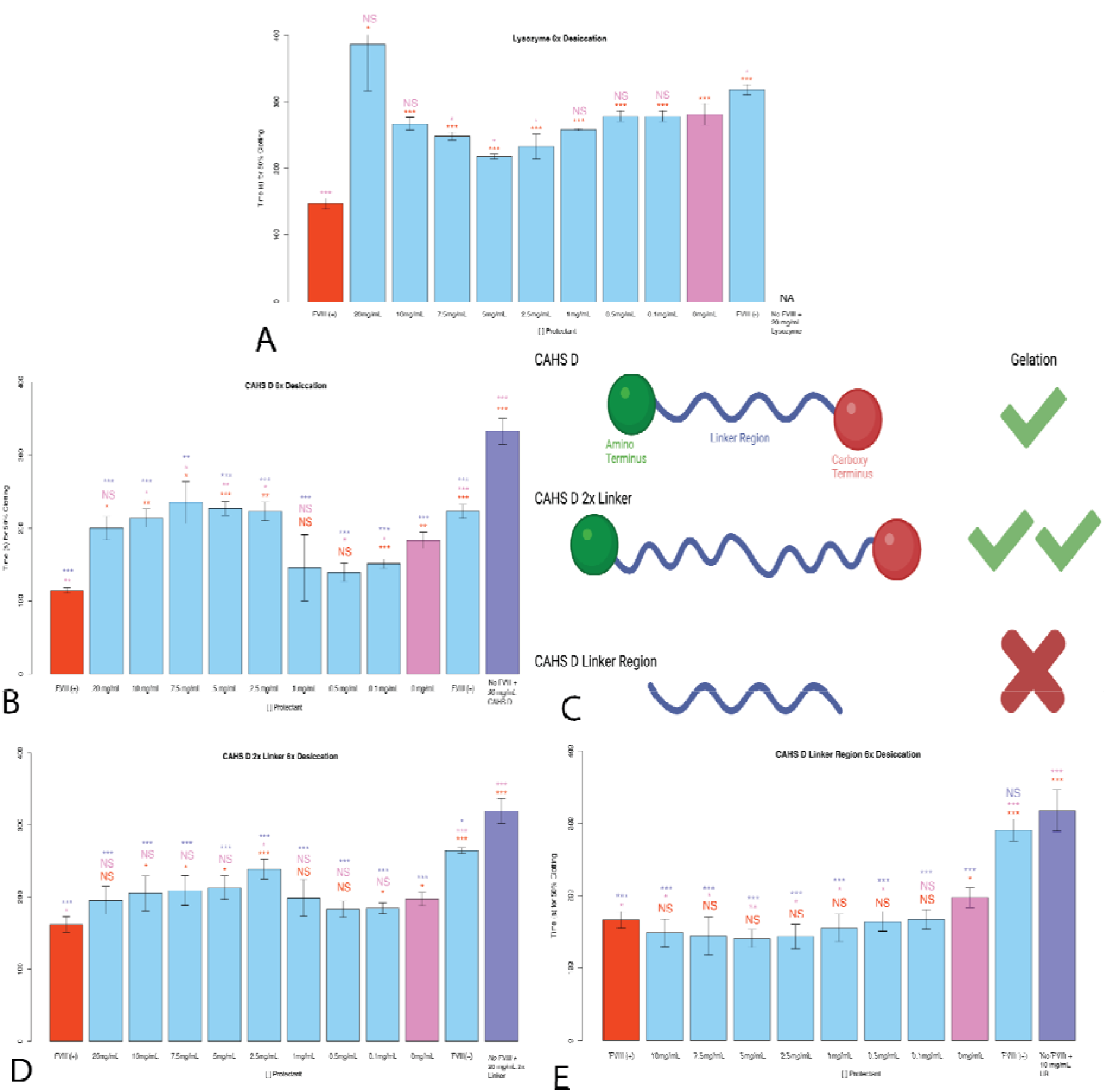
Protective capacity of natural and engineered CAHS D variants during repeated desiccation cycles. Histograms of the 50% clotting time of FVIII after various treatments with a (A) control protein lysozyme, (B) CAHS D, (D) CAHS D 2X Linker and (E) CAHS D Linker Region, subjected to repeated desiccation cycles. “FVIII (+)” represents FVIII deficient human blood plasma treated with FVIII. All red notations above sample bars represent statistical comparison with “FVIII (+).” “X mg/mL” notations beneath sample bars correspond to the concentrations of CAHS D or applicable CAHS D variant mixed with FVIII before desiccation. All pink notations above sample bars represent statistical comparison with “0 mg/mL.” “FVIII (-)” represents FVIII deficient human blood plasma treated with no FVIII. “No FVIII + X mg/mL CAHS D or CAHS D variant” represents FVIII deficient human blood plasma treated with X mg/mL CAHS D or the indicated CAHS D variant, but no FVIII. All purple notations above sample bars represent statistical comparison “No FVIII + X mg/mL CAHS D or CAHS D variant.” “NA” denotes that although the relevant experiment was conducted no clotting was observed during the entire duration of the clotting assay. Notations above sample bars represent statistical comparisons by 2-tailed T-test. Error bars represent bi-directional standard deviation. P value > 0.05 = NS, P < 0.05 = *, P < 0.01 = **, P < 0.005 = ***. (C) Schematic depiction of CAHS D, CAHS D 2X Linker, and CAHS D Linker Region protein ensembles and an indication of whether or not these proteins form hydrogels.

After observing the minimal protective capacity displayed by lysozyme, we tested FVIII mixed with CAHS D prior to drying cycles. At low concentrations (0.1-0.5 mg/mL) CAHS D confers partial protection or full protection of FVIII function during desiccation (Fig. 3B). However, beyond 1 mg/mL CAHS D became inhibitory to clotting, with 50% clotting times being longer than in 0 mg/mL samples (Fig. 3B). Consistent with this observation, FVIII DHBP treated with 20 mg/mL CAHS D clotted significantly more slowly than FVIII DHBP alone, indicating that CAHS D at high concentration interferes with human plasma clotting (Fig. 3B, purple bar). However, the observation that between 0.1-0.5 mg/mL, CAHS D-treated FVIII can still clot FVIII DHBP even after repeated desiccation cycles indicates that at low concentrations, the protective capacity of CAHS D outweighs its inhibitory clotting effects.

CAHS D is known to undergo a concentration dependent phase transition from a solution into a solid gel state^20,28,29^. At concentrations of 0.1 - 7.5 mg/mL CAHS D, the protein is diffuse in solution, however at ~10 mg/mL and higher, CAHS D forms a hydrogel^20^. We reasoned that the counterintuitive result that low concentrations, but not high concentrations, of CAHS D protect FVIII might be linked to CAHS D’s propensity to form a hydrogel^20^. This could also explain why plasma 50% clotting time was slowed at 20 mg/mL of CAHS D.

To investigate the role of CAHS D hydrogel formation in FVIII protection, we tested the protection efficacy of engineered CAHS D variants with different hydrogel formation behaviors. Gelation of CAHS D requires an amino terminus (N-term), linker region, and carboxy terminus (C-term) (Fig. 3C)^20^. Alterations to this native sequence and conformational ensemble influence CAHS D hydrogel formation, with expansion of the linker region leading to stronger hydrogel formation at lower concentrations, or conversely, disruption of the N-, linker, or C-term leading to no gel formation (Fig. 3C)^20^.

We first used an engineered CAHS D variant with an enhanced propensity for gelation, called 2X Linker^20^. The 2X Linker protein is composed of the N- and C-termini of CAHS D held apart by a tandemly duplicated linker region. This particular variant forms a gel at concentrations lower than CAHS D (5 mg/mL *versus* 10 mg/mL)^20^.

Consistent with our speculation that stronger gelation should decrease protective capacity during repeated desiccation cycles, 2X Linker’s protective capacity differed from CAHS D’s in that 2X Linker provided no protection to FVIII after desiccation at any concentrations examined (Fig. 3B&D).

Finally, we used a CAHS D variant that does not form a hydrogel at any concentration, termed CAHS D Linker Region (LR)^20^. The LR variant was constructed by removing both the N- and C-termini of CAHS D, which are required for hydrogel formation, resulting in just the linker region20. When treated with any concentration of LR above 0.1 mg/mL we observed robust and complete protection of FVIII function after desiccation (Fig. 3E).

Combined, these results suggest that the desiccation-related IDP, CAHS D, provides increased protection to FVIII subjected to repeated desiccation cycles, relative to a control protein (lysozyme). Furthermore, by engineering CAHS D such that it cannot form a hydrogel, we are able to enhance the ability of CAHS D to confer protection to FVIII during repeated desiccation cycles.

### Robust stabilization of FVIII during repeated cycles of drying and rehydration is not an ubiquitous ability of all IDPs

To investigate whether other desiccation-related IDPs beyond CAHS D and its variants are able to protect FVIII during repeated desiccation cycles, we selected and tested two additional IDPs: AavLEA1 and Hero9. AavLEA1, is a Late Embryogenesis Abundant (LEA) protein from the desiccation tolerant nematode *Aphelenchus avenae*. LEA proteins, like CAHS D, have been shown to be sufficient in protecting biomolecules from desiccation both *in vivo* and *in vitro^30–32^*, but unlike CAHS proteins are not known to form hydrogels. Hero9 belongs to a newly discovered class of proteins called Heat-resistant obscure (Hero) proteins, from the human proteome, which despite being derived from a non-desiccation tolerant organism, have been observed to confer protection against protein instability and aggregation^33,34^. Similar to LEA proteins, Hero9 has not been reported or observed to form gels.

Neither AavLEA1 nor Hero9 independently interfere with FVIII DHBP clotting (4A&B, purple bars). When Hero9 was mixed with FVIII prior to repeated desiccation cycles it was observed to confer partial protection to FVIII at intermediate concentrations (2.5-10 mg/mL; Fig. 4A). Suggesting that while Hero9 may be protective to some biomolecules under certain conditions^33,34^, this effect does not extend to repeated drying cycles with FVIII. However, contrary to our results with Hero9, AavLEA1 was observed to be fully or partially protective of FVIII after repeated desiccation at concentrations equal to or above 0.1 mg/mL (Fig. 4B). This is in line with our measurements of LR protection (Fig. 3E).

**Figure 4.**
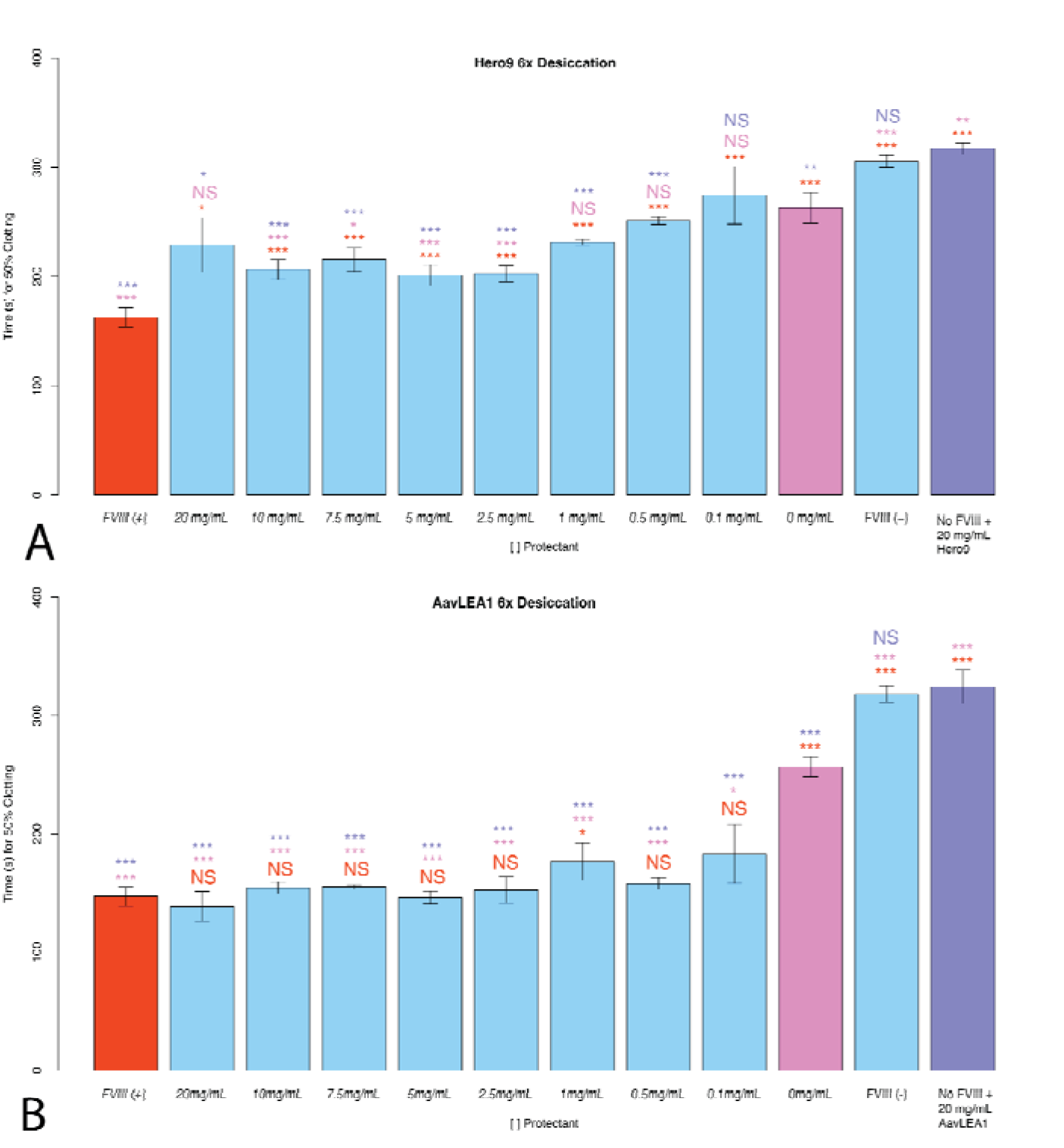
Non-CAHS IDPs’ protective capacity during repeated desiccation cycles. Histograms of the 50% clotting time of FVIII with different treatments (A) Hero9 and (B) AavLEA1. “FVIII (+)” represents FVIII deficient human blood plasma treated with FVIII. All red notations above sample bars represent statistical comparison with “FVIII (+).” “X mg/mL ‘‘ notations beneath sample bars correspond to the concentrations of AavLEA1 or Hero9, mixed with FVIII before desiccation. All pink notations above sample bars represent statistical comparison with “0 mg/mL.” “FVIII (-)” represents FVIII deficient human blood plasma treated with no FVIII. “No FVIII + X mg/mL AavLEA1 or Hero9” represents FVIII deficient human blood plasma treated with X mg/mL AavLEA1 or Hero9, but no FVIII. All purple notations above sample bars represent statistical comparison with “No FVIII+ X mg/mL, AavLEA1, or Hero9.” “NA” denotes that although the relevant experiment was conducted no clotting was observed during the entire duration of the clotting assay. Notations above sample bars represent statistical comparisons by 2-tailed T-test. Error bars represent bi-directional standard deviation. P value > 0.05 = NS, P < 0.05 = *, P < 0.01 = **, P < 0.005 = ***

Taken together, our results suggest that FVIII can be efficiently protected during repeated dehydration/rehydration cycles by both sugar and protein-based mediators of desiccation tolerance, but that robust preservation of FVIII by IDPs is not uniform and there may be sequence features as well as biochemical, and/or biophysical properties that make an IDP more or less protective for a particular client under a specific stress.

### Classic sugar-based mediators of desiccation tolerance stabilize FVIII during heat stress in a dry state

Beyond repeated rehydration/dehydration cycles, another stress potentially encountered by a biologic during storage or transportation outside of the cold-chain is thermal stress. Nonreducing disaccharides are known to improve thermotolerance of live cells and act to stabilize labile biomolecules exposed to increased temperature both *in vivo* and *in vitro^35,36^*. After establishing the protective capacity of sucrose and trehalose under repeated desiccation stress, we tested these sugars’ ability to stabilize FVIII in a dry state under heating stress. To test this, we subjected FVIII which had been dried once to 48 hours of heating at 95 °C.

Heating of dry FVIII without trehalose or sucrose resulted in total loss of the factors clotting ability (Fig. 5A&B). However, similar to 6X desiccation trials (Fig. 3A&B), sucrose and trehalose both confer significant protection to FVIII under thermal stress (Fig. 5A&B). During heat stress in a dry state, sucrose provided partial protection to FVIII at all concentrations tested (0.1-20 mg/mL; Fig. 5A), while trehalose provided partial protection at low concentrations (0.5-5 mg/mL) and full protection at higher (7.5-20 mg/mL) concentrations (Fig. 5B).

**Figure 5.**
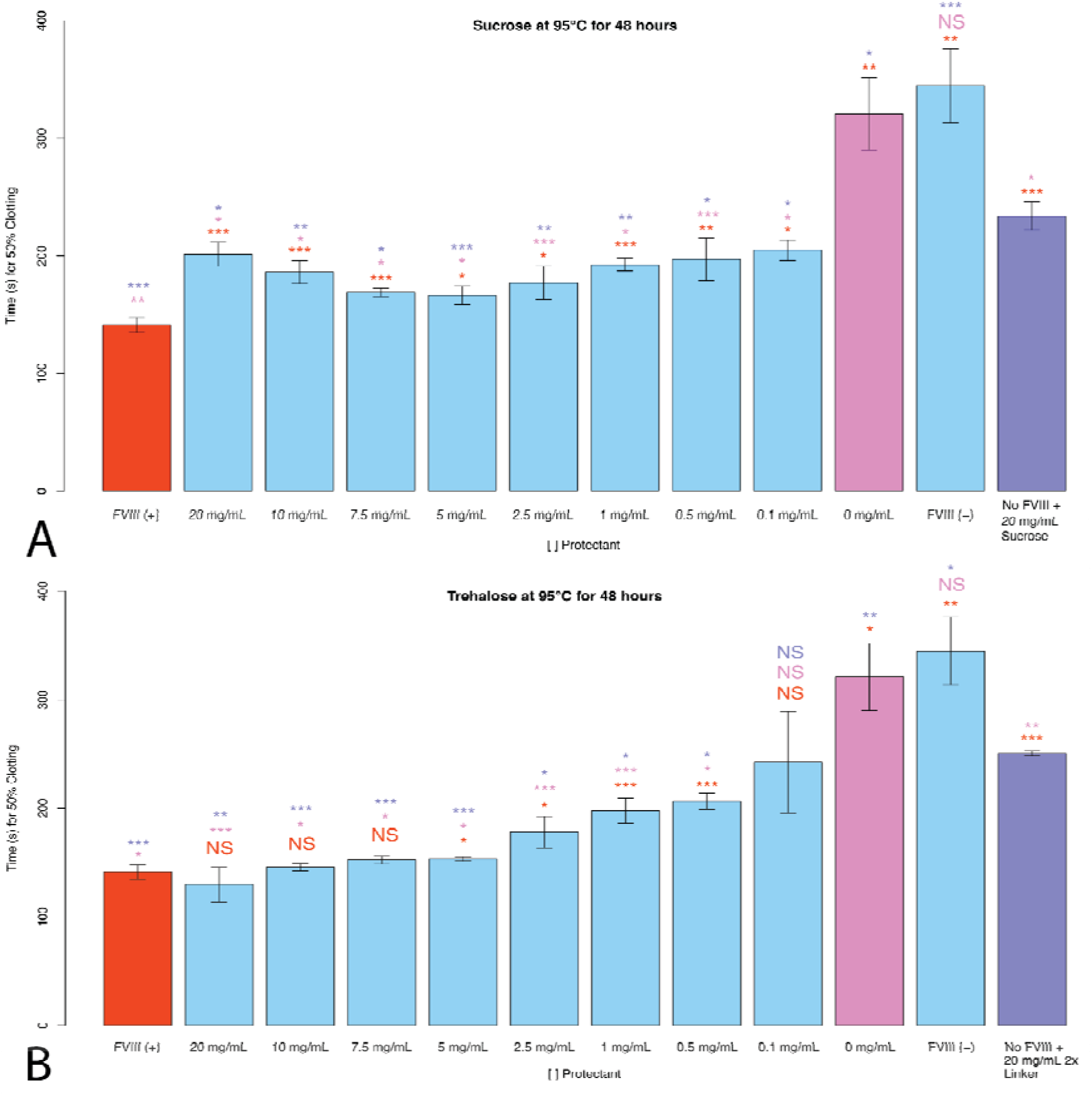
Using non-reducing disaccharides as potential stabilizers of FVIII under thermal stress. Histogram of the 50% clotting time of FVIII coincubated with (A) sucrose or (B) trehalose. “FVIII (+)” represents FVIII deficient human blood plasma treated with unstressed FVIII. All red notations above sample bars represent statistical comparison with “FVIII (+).” “X mg/mL” notations beneath sample bars correspond to the concentrations of non-reducing disaccharide mixed with FVIII before desiccation. All pink notations above sample bars represent statistical comparison with “0 mg/mL.” “FVIII (-)” represents FVIII deficient human blood plasma without FVIII supplementation. “No FVIII + X mg/mL Sugar ‘‘ represents FVIII deficient human blood plasma treated with X mg/mL non-reducing disaccharide, but no FVIII. All purple notations above sample bars represent statistical comparison with “No FVIII + X mg/mL Sugar” sample bar. Notations above sample bars represent statistical comparisons by 2-tailed T-test. Error bars represent bi-directional Standard Deviation. P value > 0.05 = NS, P < 0.05 = *, P < 0.01 = **, P < 0.005 = ***

These results demonstrate that the thermal protection observed of sugar-based protectants for model enzymes extends to the thermal protection of the biologic FVIII.

### CAHS D and engineered CAHS D variants stabilize FVIII during heat stress in a dry state

CAHS proteins, including CAHS D, are known to be heat soluble in solution^37^ and they have been shown to increase thermal tolerance when heterologously expressed in dry yeast^12^, but their ability to confer long-term thermal tolerance to a client has yet to be established *in vitro*.

As a negative control we tested lysozyme’s ability to stabilize FVIII in a dry state under thermal stress (95 °C for 48 hours). Similar to the results from repeated desiccation cycles, lysozyme was not protective to FVIII under heating stress at any concentration (Fig. 6A). Furthermore, high concentrations (15-20 mg/mL) of lysozyme mixed with FVIII inhibited blood clotting all together (Fig. 6A). This complete inhibition was not observed for lysozyme in experiments conducted with repeated desiccation cycles (Fig. 3A), suggesting that heating may impart some detrimental change to heat-insoluble lysozyme which interferes with plasma clotting.

**Figure 6.**
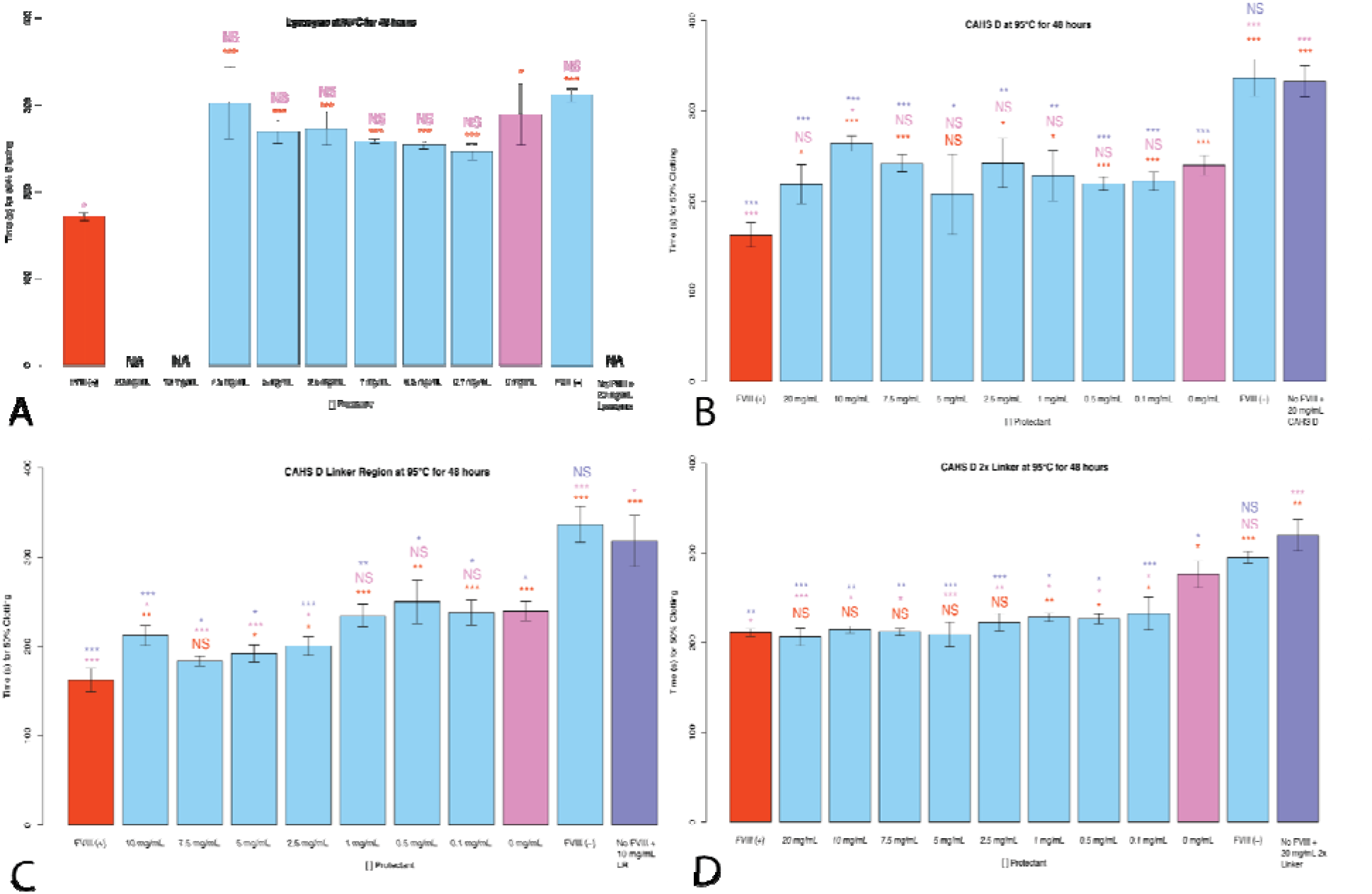
Using CAHS D and engineered CAHS D variants as potential stabilizers of FVIII under thermal stress. Histograms of 50% clotting time of FVIII treated with (A) lysozyme, (B) CAHS D, (C) CAHS D Linker Region, or (D) CAHS D 2X Linker prior to desiccation and thermal stress. “FVIII (+)” represents FVIII deficient human blood plasma treated with nonperturbed FVIII. All red notations above sample bars represent statistical comparison with “FVIII (+).” “X mg/mL’’ notation beneath sample bars correspond to the concentration of CAHS D or the indicated CAHS D variant mixed with FVIII before desiccation. All pink notations above sample bars represent statistical comparison with “0 mg/mL.” “FVIII (-)” represents FVIII deficient human blood plasma not treated with FVIII. “No FVIII + X mg/mL CAHS D or CAHS D variant” represents FVIII deficient human blood plasma treated with X mg/mL CAHS D or CAHS D variant, but no FVIII. All purple notations above sample bars represent statistical comparison with “No FVIII + X mg/mL CAHS D or CAHS D variant”. “NA” denotes that, although the relevant experiment was conducted, no clotting was observed during the entire duration of the clotting assay. Notations above sample bars represent statistical comparisons by 2-tailed T-test. Error bars represent bi-directional standard deviation. P value > 0.05 = NS, P < 0.05 = *, P < 0.01 = **, P < 0.005 = ***

Coincubation of FVIII with heat soluble CAHS D during heat stress showed no protection at any concentration and in one case (10 mg/mL) showed antagonism (Fig. 6B). Similar to our 6X desiccation experiments, LR showed improved protection relative to CAHS D, with partial or full protection after heat stress observed at high concentrations (7.5-20 mg/mL) (Fig. 6C). Interestingly, when we tested the 2X Linker variant under thermal stress we observed robust protection of FVIII (Fig. 6D). The 2X Linker construct provided partial protection at low concentrations (0.1-1 mg/mL) and full protection at all concentrations above 1 mg/mL (Fig. 6D).

These results suggest that sequence features as well as biophysical and biochemical properties can be tuned to make IDP-based protectants more or less effective at preventing damage from specific types of stress.

### Stabilization of FVIII during heat stress in a dry state is not a ubiquitous ability of all heat soluble IDPs

Like CAHS proteins, LEA and Hero proteins are known to be heat soluble^12,33,37,38^. We wondered if heat solubility of IDPs might be a feature that commonly confers thermal tolerance to client proteins. To test this we incubated FVIII with AavLEA1 or Hero9 prior to desiccation and heating for 48 hours at 95 °C.

After mixing AavLEA1 with FVIII and subjecting dry samples to thermal stress, FVIII function was observed to have no protection under heat stress at low to medium concentrations (0.1-2.5 mg/mL) and completely preserved at high concentrations (5-20 mg/mL) (Fig. 7A). Conversely, after heating, Hero9 did not confer protection to FVIII at any concentrations, and actually became antagonistic, causing slowed 50% clotting time, at high concentrations (7.5-20 mg/mL) (Fig. 7B).

**Figure 7.**
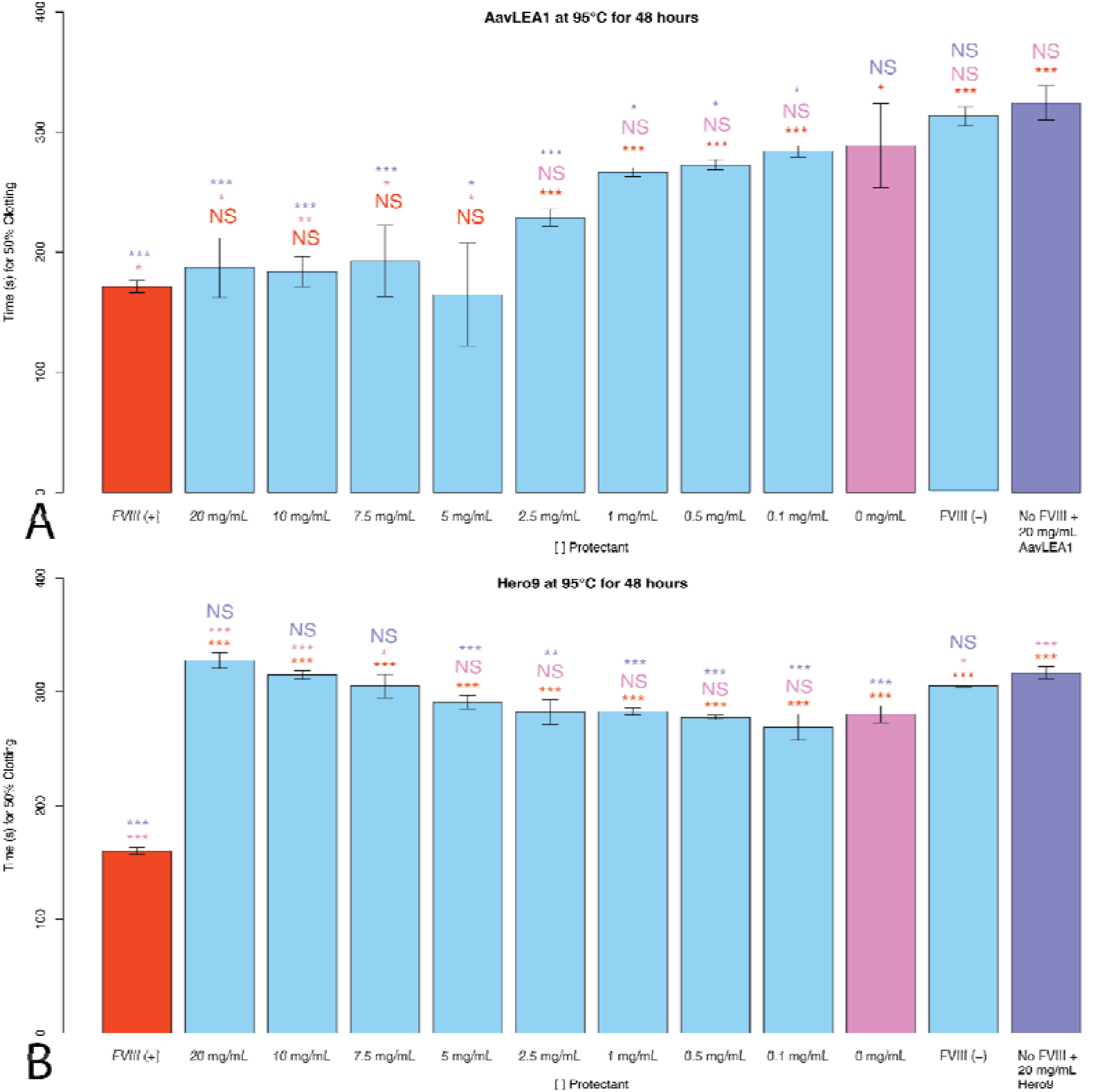
Using AavLEA1 or Hero9 as potential stabilizers of FVIII under thermal stress. Histogram representation of the 50% clotting time of FVIII treated with (A) AavLEA1, (B) Hero9. “FVIII (+)” represents FVIII deficient human blood plasma treated with unperturbed FVIII. All red notations above sample bars represent statistical comparison with “FVIII (+).” “X mg/mL” notation beneath sample bars correspond to the concentrations of AavLEA1 or Hero9 mixed with FVIII before desiccation. All pink notations above sample bars represent statistical comparison with “0 mg/mL.” “FVIII (-)” represents FVIII deficient human blood plasma treated with no FVIII. “No FVIII + X mg/mL AavLEA1 or Hero9” represents FVIII deficient human blood plasma treated with X mg/mL AavLEA1 or Hero9 but no FVIII. All purple notations above sample bars represent statistical comparison with “No FVIII+ X mg/mL, AavLEA1, or Hero9.” Notations above sample bars represent statistical comparisons by 2-tailed T-test. Error bars represent bi-directional standard deviation. P value > 0.05 = NS, P < 0.05 = *, P < 0.01 = **, P
< 0.005 = ***

Taken together, these data indicate that while other desiccation-related and heat soluble IDPs can protect FVIII against thermal stress, this is not a ubiquitous feature of all heat soluble IDPs.

### CAHS D LR is able to stabilize FVIII in a dry state for at least 10 weeks

Considering the practical application of using dry preservation, or xeroprotection, as a means of stabilization of labile biologics during their transport and storage, we wondered how shelf-stable FVIII might be in a dry state.

To test this, we selected our LR variant since this variant showed the best protection under repeated drying cycles (Fig. 3E). FVIII was incubated with or without the highest concentration of LR used in other experiments, 10 mg/mL. Samples were split, with one sample being left hydrated while the other was dried. Triplicate samples were prepared and examined every week for ten weeks (Fig. 8). FVIII left in a hydrated state with or without LR lost a significant amount of functionality over the course of 10 weeks (p < 0.005; Fig. 8). Dried samples without LR lost less clotting functionality compared to hydrated samples, but still lost a significant amount of clotting ability over the course of ten weeks (p < 0.005; Fig. 8). Samples of FVIII dried with 10 mg/mL of LR did not statistically change over the course of 10 weeks (Fig. 8).

**Figure 8.**
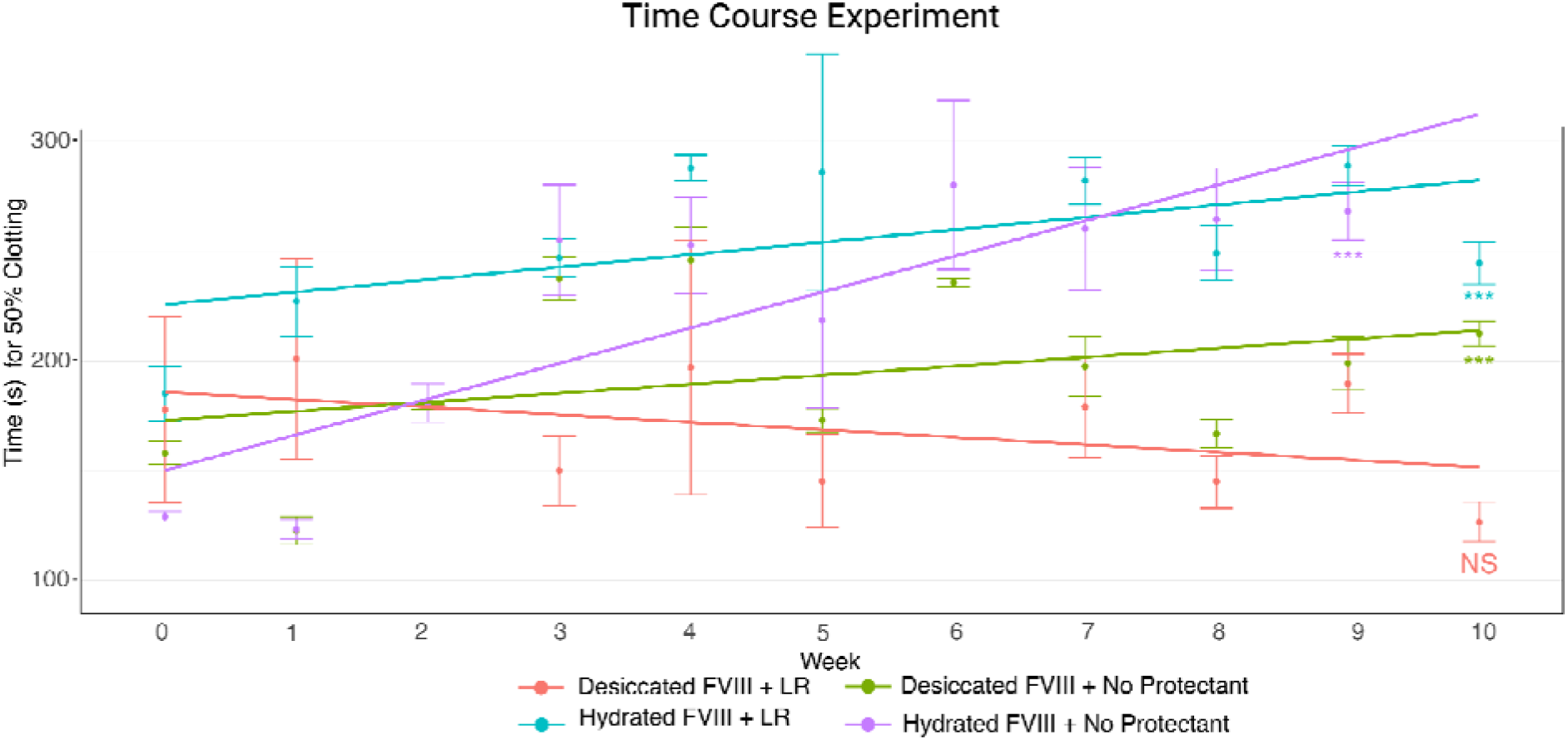
CAHS D Linker Region time course stabilization. 50% clotting time of FVIII in a hydrated or dry state with or without addition of 10 mg/mL of CAHS D Linker Region. Samples were prepared and left dry/hydrated for 1 to 10 weeks before testing in our clotting assay. Notations below sample bars represent statistical comparisons between the first and the last time point taken using a 2-tailed T-test. Error bars represent bi-directional standard deviation. P value > 0.05 = NS, P < 0.05 = *, P < 0.01 = **, P < 0.005 = ***

These data indicate that while FVIII clotting potential decreases over time when stored under hydrated conditions (with or without LR) or when stored dry without an excipient, the addition of our engineered LR protein at high concentration (10 mg/mL) is sufficient to stabilize FVIII function for at least 10 weeks.

## Discussion and conclusions

Here we demonstrate that dry preservation of Human Blood Clotting Factor VIII, a key molecule in the intrinsic blood clotting pathway with numerous clinical applications, is achievable using both sugar and protein-based excipients.

It is of interest to note that while FVIII can be stabilized under repeated desiccation cycles and thermal stress, the mediators that do best at preventing damage during these distinct stresses are different. The non-gelling protein CAHS D Linker Region performed best at preventing damage to FVIII in repeated desiccation cycles, while the gelling variant CAHS D 2X Linker outperformed all other excipients in preserving FVIII function under thermal stress. These results suggest that, using classic protein engineering approaches, stress-related IDPs can be tuned to serve specific functions with regard to biologic stabilization.

Beyond classic sequence-based engineering approaches, the use of IDPs offers other advantages not fully explored in this work, such as the ability to augment functions and protective mechanisms using different chemical environments. IDPs are, by their nature, highly sensitive to their chemical environment given their relative lack of intramolecular bonds and high solvent exposed surface area^40,41^. In line with this, recently, it has been shown that CAHS D’s function is highly influenced by its chemical environment^19^. Specifically, CAHS D works synergistically with trehalose to promote desiccation tolerance both *in vitro* and *in vivo^19^*. It is of note that CAHS D is naturally enriched alongside trehalose as tardigrades desiccate and this synergistic effect was not as pronounced for CAHS D mixed with another, non-tardigrade sugar, sucrose^19^. This suggests that not only can IDP protective function be tuned by changes to their sequence, but also by modulation of their chemical environment.

We also observed that biopreservation properties do not extend to all proteins (e.g., lysozyme) nor even all IDPs (e.g., Hero9). Deciphering the exact molecular grammar that makes an IDP protective or not during desiccation/thermal stress will contribute to the continued development methods for xeroprotection of biologics.

This study serves as a proof of principle that dry preservation methods can be effective in protecting labile biologics, offering a convenient, logistically simple, and economically viable means of stabilizing life saving medicines. This will not only be beneficial for global health initiatives in remote or developing parts of the world, but also for fostering a safe and productive space economy which will be reliant on new technologies that break our dependance on the coldchain for the storage of medicine, food, and other biomolecules.

## Materials and methods

### Cloning and expression of CAHS D, CAHS D variants, AavLEA1, and Hero9

Constructs were cloned via Gibson Assembly in pET28b and transformed into BL21 (DE3) *E. coli* (New England Biolabs). Transformed *E. coli* were then plated on Luria-Bertani (LB) agar with 50□μg/mL Kanamycin (Kan). Constructs were expressed in 2.8 L flasks containing 1 L of LB and 50 μg/mL Kan. Inoculated media was shaken at 37 °C, 180 rpm (Eppendorf Innova S44i) until an optical density (OD) of 0.6 was reached. Expression was induced with 1 mM IPTG for 4 hours. Cells were harvested by centrifugation at 3,500 rpm for 30 minutes at 4 °C. Cells were resuspended in 5 mL of pellet resuspension buffer (20 mM Tris pH 7.5) supplemented with protease inhibitors. Pellets were stored at −80 °C until use.

### Protein purification of CAHS D, CAHS D variants, and AavLEA1

Pellets were thawed at room temperature and subjected to heat lysis in boiling water for 10 minutes and allowed to cool for 15 minutes. Samples were then centrifuged at 10,500 rpm for 45 minutes at 10 °C. The supernatant was filter-sterilized through a 0.22 μm syringe filter (EZFlow Syringe Filter, Cat. 388-3416-OEM) to remove any insoluble particles. The filtrate was diluted two times the volume with buffer UA (8 M Urea, 50 mM sodium acetate, pH 4). This was loaded onto a HiPrep SP HP 16/10 cation exchange column (Cytiva) and purified on an AKTA Pure, controlled using the UNICORN 7 Workstation pure-BP-exp.

AavLEA1 was eluted using 0-70% UB gradient (8 M Urea, 50 mM sodium acetate, and 1 M NaCl, pH 4) and fractionated over 15 column volumes. CAHS D, CAHS D 2X Linker and CAHS D Linker Region were eluted using a 0-40% UB gradient and were fractionated over 15 column volumes. Purified protein fractions were confirmed using SDS-PAGE and selected fractions were pooled for dialysis in 3.5 kDa tubing in 20 mM sodium phosphate buffer pH 7. This was followed by six rounds of dialysis in Milli-Q water (18.2 MΩcm) at 4 hour intervals each. Samples were quantified fluorometrically (Invitrogen Qubit 4 Fluorometer, REF Q33226), flash frozen, then lyophilized for 48 hours (Labconco FreeZone 6, Cat. 7752021) and stored at −20 °C until further use.

### Protein purification of Hero9

Pellets were thawed at room temperature and subjected to heat lysis in boiling water for 10 minutes and allowed to cool for 15 minutes. The pellets were further lysed by sonication for 5 minutes, using 40% amplitude and 30 seconds on - 30 seconds off cycles on ice. All insoluble components were removed via centrifugation at 10,500 rpm at 10 °C for 30 minutes. The supernatant was filtered with 0.45 μm and 0.22 μm syringe filters (EZFlow Syringe Filter, Cat. 388-3416-OEM). The protein was then purified using an anion exchange HiPrep Q HP 16/10 (Cytiva, Cat. 29018183) on the AKTA Pure 25 L (Cytiva, Cat. #29018224), controlled using the UNICORN 7 Workstation pure-BP-exp. Protein was eluted using a gradient of 0-70% B (25 mM Tris-HCl, 1 M NaCl, pH 7.4), over 20 column volumes. Fractions were assessed by SDS-PAGE and pooled for dialysis in 3.5 kDa MWCO dialysis tubing (SpectraPor 3 Dialysis Membrane, Part No. 132724). Protein was dialyzed at 25 °C for 4 hours against 25 mM Tris-HCl, 150 mM NaCl, pH 7.4, then transferred to 25 mM Tris-HCl, 50 mM NaCl, pH 7.4 overnight. This was followed by 4 rounds of 4 hours each in Milli-Q water (18.2 MΩcm). Dialyzed samples were quantified fluorometrically (Invitrogen Qubit 4 Fluorometer, REF Q33226), aliquoted in the quantity needed for each assay, lyophilized (Labconco FreeZone 6, Cat. 7752021) for 48 hours, then stored at −20 °C until use.

### Formulation of FVIII and protectants

FVIII was obtained in a lyophilized state (Sigma Aldrich cat. H0920000-3EA), and rehydrated using molecular grade water. Throughout this research, multiple different FVIII vials were used, all originating from the same lot (lot 6). Lyophilized FVIII deficient human blood plasma (Helena Biosciences cat. 5193) was rehydrated in accordance with Helena Biosciences blood plasma protocol. Over the course of the experiments, multiple FVIII deficient human blood plasma vials were combined and mixed to mitigate potential 50% clotting time discrepancies within each vial/batch. All FVIII deficient plasma in this study originated from the same lot (lot 2-22-5193).

### FVIII Clotting Assay

Pacific Hemostasis’s standardized coagulation APTT-XL protocol was followed, using FVIII as a model clotting factor in this assay^42^. To test the clotting function, 50 μL FVIII deficient human blood plasma was treated with 5 μL of 5 mg/mL (25 μg) FVIII. After addition of FVIII into the FVIII deficient human blood plasma, 50 μL of APPT-XL reagent was added to the mixture and incubated at 37 □ for 3 minutes. After incubation, 50 μL of CaCl pre-heated to 42 □ was added. FVIII function was determined by 50% clotting time (aPTT). aPTT was measured using a platereader (Tecan Spark600 Cyto) to record the absorbance at 405 nm every 15 seconds over a 7 minute period^42^.

### 6X desiccation protocol

10 μL of 5 mg/mL (50 μg) FVIII was desiccated for 1 hour in a SpeedVac (Savant SpeedVac SC110 Vacuum Concentrator Model SC110-120) and rehydrated with 10 μL molecular grade water. This cycle of desiccation and rehydration was repeated 6 times for each 6X desiccation treated sample. After the final hour of desiccation, the FVIII was rehydrated in 10 μL of molecular grade water. 5 μL (25 μg) of treated FVIII was added to FVIII deficient human blood plasma to quantify subsequent clotting activity. To test the potential protective effects of sugar/protein excipients, 5 μL of 10 mg/mL (50 μg) FVIII was incubated with 5 μL sugar/protein excipients at varying concentrations for a final FVIII concentration of 5 mg/mL FVIII prior to being subjected to the 6X desiccation protocol. Following 6X desiccation, aPTT was tested on control and experimental samples using the FVIII clotting assay outlined above.

### Desiccation and heat stress protocol

In order to establish a baseline for heat stress, 10 μL of 5 mg/mL (50 μg) FVIII was desiccated for 1 hour in a SpeedVac (Savant SpeedVac SC110 Vacuum Concentrator Model SC110-120) and then placed at 95 □ for 48 hours. After 48 hours of heating, the FVIII was rehydrated in 10 μL molecular grade water. 5 μL (25 μg) of treated FVIII was added to FVIII deficient human blood plasma to quantify subsequent clotting activity. To test the potential protective effects of sugar/protein excipients, 5 μL of 10 mg/mL (50 μg) FVIII was treated and incubated with 5 μL sugar/protein excipients at varying concentrations for a final FVIII concentration of 5 mg/mL FVIII prior to being subjected to the desiccation and heating protocol. Following desiccation and heating, control and experimental samples were tested for aPTT using the FVIII clotting assay outlined above.

### Time course

FVIII function was measured over a period of 10 weeks. 10 samples in triplicate (one sample for each week) consisting of 5 μL of 10 mg/mL FVIII with 5 μL of 20 mg/mL CAHS D Linker Region resulting in 50 μg of FVIII and 10 mg/mL CAHSD Linker Region, were desiccated for 1 hour. Conversely, 10 μL of 5 mg/mL (50 μg) FVIII without any excipient was desiccated for 1 hour. All dried samples were then placed in 1.5 mL Eppendorf tubes sealed with parafilm. The tubes were left on the benchtop at ambient room temperature, ~22 □. At one week intervals, a sample would be rehydrated with 10 μL molecular grade water. The 1st sample set of each experiment being measured 1 week after initial desiccation and so on. 5 μL (25 μg) FVIII would be withdrawn from each sample and used to determine aPTT using the FVIII clotting assay outlined above.

For the hydrated FVIII samples, one sample was prepared in triplicate for hydrated FVIII with no protectant and hydrated FVIII supplemented with 10 mg/mL CAHS D Linker Region. Each replicate of hydrated FVIII with no protectant sample contained 5 mg/mL FVIII in molecular grade water. Each replicate of hydrated FVIII with CAHS D Linker Region contained 5 mg/mL FVIII resuspended in molecular grade water with CAHS D Linker Region at a final concentration of 10 mg/mL. All hydrated samples were placed in 1.5 mL Eppendorf tubes sealed with parafilm. The tubes were left on the benchtop at ambient room temperature, ~22 □. At one week intervals for 10 weeks, 5 μL (25 μg) FVIII would be withdrawn from each sample and used to determine a aPTT using the FVIII clotting assay outlined above.

### Statistics

For each clotting experiment, the time for 50% clotting was determined. Using this value, standard deviation of the triplicate within each trial is represented by two sided error bars. Significance values are attained by conducting a two-sided T-Test. Significance was noted for P-values less than or equal to 0.05 as noted in figure legends.

## Acknowledgments

This work was supported by DARPA (W922NF-20-2-0137) to TCB. In addition, this work was made possible in part through support from an Institutional Development Award (IDeA) from the National Institute of General Medical Sciences of the National Institutes of Health (Grant # 2P20GM103432). MHP was supported in part by the Wyoming NASA Space Grant Consortium, NASA Grant #80NSSC20M0113, INBRE undergraduate research grant through the Institutional Development Award (IDeA) from the National Institute of General Medical Sciences of the National Institutes of Health under Grant #2P20GM103432, and the Wyoming Research Scholars Program. The authors are thankful to members of the Boothby Lab for discussions and reading of this manuscript. We thank members of the Water and Life Interface Institute (WALII), supported by NSF DBI grant #2213983, for helpful discussions.

## Author contributions

MHP, SS, and TCB participated in experimental design, data analysis and preparation of the manuscript and figures. MHP and SS performed clotting assays. MHP, SS, SB, SK, KN participated in protectant expression and purification. The authors have read and approved the final manuscript.

## Materials and Correspondence

Correspondence and material requests should be made to Thomas Boothby.

